# The evolution of Monkeypox virus: a genetic and structural analysis reveals mutations in proteins involved in host-pathogen interaction

**DOI:** 10.1101/2022.06.22.497195

**Authors:** Domenico Benvenuto, Serena Vita, Stefano Pascarella, Martina Bianchi, Marta Giovanetti, Roberto Cauda, Emanuele Nicastri, Antonio Cassone, Massimo Ciccozzi

**Author notes:** **Correspondence to:** Dr. Domenico Benvenuto, MD, Unit of Medical Statistics and Molecular Epidemiology, University Campus Bio-Medico of Rome, Via Álvaro del Portillo, 21, 00128, Rome, Italy.

## Abstract

**Background:** Over the past few months, we have witnessed a new outbreak of a MPXV that has been detected without a clear link to Africa and has quickly spread globally.

**Methods:** In this article we investigate the mutational pattern of the MPXV and provide evidence for the presence of 6 new mutations that appear to characterize the current MPX-2022 outbreak. With the use of a number of chemical and physical parameters, we predict the stability of the mutated proteins, and propose an interpretation of the impact of these mutations on viral fitness).

**Findings:** Most mutations, particularly the Immunomodulator A46, TNFr and Large transcript constituent, affect proteins playing an important role in host response to MPVX infection and could also be relevant to the clinical features of the 2022 MPXV outbreak.

**Interpretation:** Although further, experimental work is necessary for a full understanding of the impact of the mutations here reported on virus replication pathways and host immunomodulation, our in-silico data suggest the importance of monitoring the emergence of new MPXV mutations for the prevention of future outbreaks potentially dangerous for public health.

**Funding:** No funding to declare

## Introduction

Human monkeypox virus (MPXV) is a double-stranded DNA virus of the Orthopoxvirus genus, family Poxviridae. The MPXV was discovered in 1958 during an outbreak in an animal facility in Copenhagen, Denmark (1). The first human case was diagnosed in 1970 in a 9-month-old baby boy in Zaire (now the Democratic Republic of the Congo, DRC) (2), and the infection has since been reported in a number of central and western African countries. Two MPXV genetic clades have been characterized: West African and Central African (Congo Basin).

The first monkeypox cases outside of Africa were reported in 2003 in the US (3) when an outbreak occurred following importation of rodents from Africa, with all people infected becoming ill after contact with pet prairie dogs. There are mounting concerns about the geographical spread and further resurgence of monkeypox. Over the past 5 decades, monkeypox outbreaks have been reported in 10 African countries and 4 countries outside Africa (4). As of 15 June, 1158 and 1882 monkeypox cases have been reported from 12 EU/EEA and seven non-EU/EEA countries, respectively. In total, 219 confirmed cases have been reported worldwide from countries where the disease is not considered to be endemic. (5) To date, all cases confirmed by PCR have been caused by the West African clade (6) that is considered to be less pathogenic than Congo Basin clade (7). Human-to-human transmission of monkeypox is well described, including nosocomial and household transmission (8,9). However, human-to-human chains of transmission have historically been less well recognized especially in West African Clade (10). The evolution of short-read, high-throughput sequencing technologies has made genomic surveys of large populations readily attainable (11).

We have here examined all available MPXV DNA sequences with the aim to shed light regarding the mutational pattern mapping of the newly West Africa genome and the patterns of selection, if any, of viral protein genes. In particular, here we describe the presence of six unique mutations present in all MPVX isolates during the 2022, the majority of which have occurred in proteins potentially relevant in MPVX-host relationship.

## Materials and Methods

A database including 300 MPXV genome sequences from NCBI database (https://www.ncbi.nlm.nih.gov/nuccore) has been built, subsequently ambiguous sequences showing low coverage or expressing non-coding regions have been removed. Sequence has been aligned using MAFFT and has been manually edited using Aliview according to the consensus Monkeypox coding reading frame. MEGA-X has been used to perform phylogenetic analysis; the evolutionary history has been inferred using the Minimum Evolution method using 1000 replicate [12]. The evolutionary distances were computed using the Tamura 3-parameter method [13].

The 3-Dimensional structure of the mRNA Capping, Ubiquitine, Poxin-Schlafen, RNA polymerase, immunomodulator A46, Transcript, Chemokine inhibitor, TNF receptor and Ankyrin has been generated using homology modeling performed with SwissModel web-based server https://swissmodel.expasy.org/interactive) while TMHMMPred v2.0 and Protter (15) have been used to evaluate the position of the protein within the membrane. CUPSAT (16) and Dynamut2 (17) online servers have been used to estimate the stability of potential mutations.

## Results

A database including 106 high coverage MPXV sequences with a total of 196694 base pairs has been built. All the sequences were aligned and manually edited, subsequently all the missense mutations were analyzed in several proteins. The minimum evolution phylogenetic analysis (Figure 1) has shown the presence of two main clades; the clade I included sequences of Western African Monkeypox strain while the Clade II included sequences of Central African Monkeypox Strain. The MPXV isolated from the recent (1022) MPVX outbreaks were grouped in the same cluster as those isolated from Israel and Singapore during 2018 and 2019, respectively.

**Figure 1.**
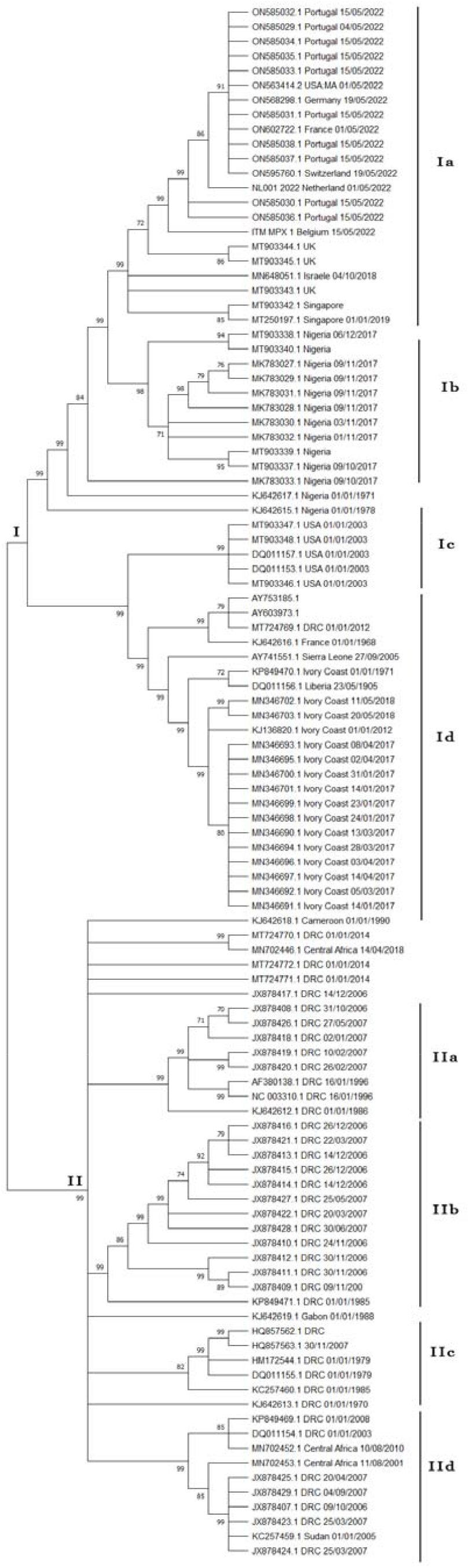
Minimum evolutionary phylogenetic tree of MPX virus

The best resulting protein template for the homology modeling, the percentage of sequence identity, the quality score Global Model Quality Estimate G (GMQE) and the quality score QMEANDisCo that provide an estimation of the global quality of the model are reported in table 1. Moreover, structural analysis has shown that none of the analyzed proteins have a potential transmembrane amino acidic region.

**Table 1.** homology modelling reporting template, Seq Identity, GMQE and QMEANDisCo Global scores

|  | Template | Seq Identity | Coverage | GMQE | QMEANDisCo Global |
| --- | --- | --- | --- | --- | --- |
| mRNA Capping | 6rfl.1.H | 98.82% | 100% | 0.91 | 0.88 ± 0.05 |
| Ubiquitine | 6y22.1.A | 25.40% | 26% | 0.11 | 0.46 ± 0.11 |
| Poxin-Schlafen | 7eg0.1.B | 91.67% | 43% | 0.35 | 0.85 ± 0.06 |
|  | 7lre.1.A | 33.33% | 57% | 0.48 | 0.63 ± 0.05 |
| RNA polymerase | 6rid.1.B | 98.88% | 100% | 0.87 | 0.88 ± 0.05 |
| immunomodulator A46 | 4lqk.1.A | 95.10% | 60% | 0.45 | 0.86 ± 0.05 |
|  | 5ezu.1.A | 93.98% | 35% | 0.19 | 0.67 ± 0.05 |
| Large Transcript protein | 1c9b.3.D | 12.90% | 33% | 0.14 | 0.39 ± 0.07 |
| Chemokine inhibitor | 2grk.1.A | 86.10% | 91% | 0.80 | 0.84 ± 0.06 |
| TNF receptor | 3alq.1.E | 43.64% | 47% | 0.34 | 0.62 ± 0.07 |
|  | 3on9.1.A | 51.57% | 46% | 0.36 | 0.75 ± 0.07 |
| Ankyrin | 5y4f.1.A | 19.59% | 64% | 0.44 | 0.59 ± 0.05 |

During the manual editing phase, the sequences have been compared with the reference MPVX West Africa sequences. We have found in all the MPXV isolates during the 2022 the presence of 5 missense mutations that were not previously reported in any other MPXV sequences, and one mutation only reported in sequences isolated in Israel and Singapore during 2018 and 2019 respectively. The mutated proteins are listed in Table 2-Moreover, we have found that sequences of MPXV isolated during an outbreak in 2017 also have typical missense mutations that were not present in the reference WA strain (Ankyrin: E176A and N552S; mRNA capping protein: A3T and A543T; RNA polymerase protein I909T; Ankyrin like protein: E176A and N552S) and shared some amino acidic change typical of the CA strain (Ubiquitine: V38I and K230R; Large Transcript protein 3: N100D; Immunomodulator A46: D24N). We have also identified the missense mutations that characterize CA strain compared to WA strain (Ubiquitine: S219L; RNA polymerase: D6V and N282D; Poxin-Schlafen protein V265I and S419R; mRNA capping protein: V537I; Large Transcript protein 3: D256E; IL-1: L227S; TNFr: A121S; Ankyrin like protein: A191V and D491N). Tyrosine Kinase, Profilin e CU-ZN peroxidase have not shown evidence of missense mutations.

**Table 2.**
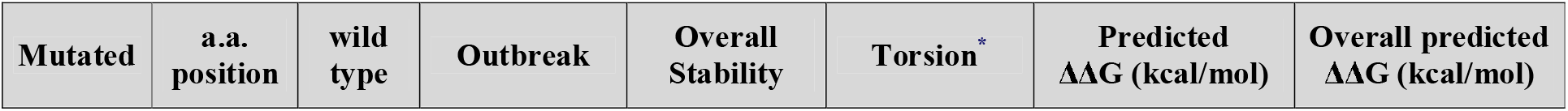
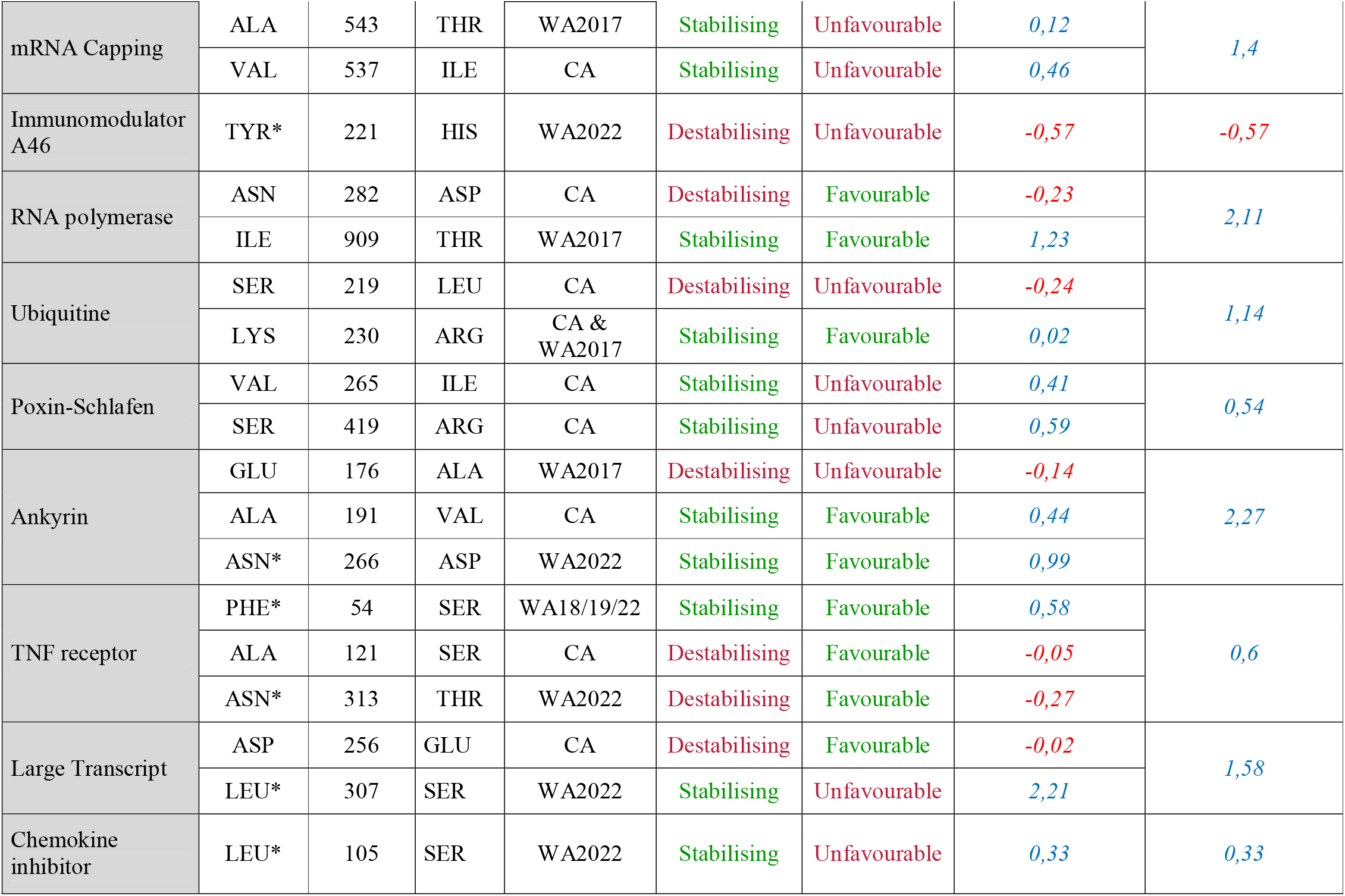
Mutational analysis reporting Mutated amino acid, amino acidic position, wild type amino acid, stability, Torsion, predicted ΔΔG (kcal/mol) and overall predicted ΔΔG (kcal/mol) * Unique mutation of 2022 outbreak

| Mutated | a.a. position | wild type | Outbreak | Overall Stability | Torsion* | Predicted $\Delta\Delta G$ (kcal/mol) | Overall predicted $\Delta\Delta G$ (kcal/mol) |
| --- | --- | --- | --- | --- | --- | --- | --- |

**Table 2** Mutational analysis reporting Mutated amino acid, amino acidic position, wild type amino acid, stability, Torsion, predicted $\Delta\Delta G$ (kcal/mol) and overall predicted $\Delta\Delta G$ (kcal/mol)
|  |  |  |  |  |  |  |  |  |
| --- | --- | --- | --- | --- | --- | --- | --- | --- |
| mRNA Capping | ALA | 543 | THR | WA2017 | Stabilising | Unfavourable | 0,12 | 1,4 |
|  | VAL | 537 | ILE | CA | Stabilising | Unfavourable | 0,46 |  |
| Immunomodulator A46 | TYR* | 221 | HIS | WA2022 | Destabilising | Unfavourable | -0,57 | -0,57 |
| RNA polymerase | ASN | 282 | ASP | CA | Destabilising | Favourable | -0,23 | 2,11 |
|  | ILE | 909 | THR | WA2017 | Stabilising | Favourable | 1,23 |  |
| Ubiquitine | SER | 219 | LEU | CA | Destabilising | Unfavourable | -0,24 | 1,14 |
|  | LYS | 230 | ARG | CA & WA2017 | Stabilising | Favourable | 0,02 |  |
| Poxin-Schlafen | VAL | 265 | ILE | CA | Stabilising | Unfavourable | 0,41 | 0,54 |
|  | SER | 419 | ARG | CA | Stabilising | Unfavourable | 0,59 |  |
| Ankyrin | GLU | 176 | ALA | WA2017 | Destabilising | Unfavourable | -0,14 | 2,27 |
|  | ALA | 191 | VAL | CA | Stabilising | Favourable | 0,44 |  |
|  | ASN* | 266 | ASP | WA2022 | Stabilising | Favourable | 0,99 |  |
| TNF receptor | PHE* | 54 | SER | WA18/19/22 | Stabilising | Favourable | 0,58 | 0,6 |
|  | ALA | 121 | SER | CA | Destabilising | Favourable | -0,05 |  |
|  | ASN* | 313 | THR | WA2022 | Destabilising | Favourable | -0,27 |  |
| Large Transcript | ASP | 256 | GLU | CA | Destabilising | Favourable | -0,02 | 1,58 |
|  | LEU* | 307 | SER | WA2022 | Stabilising | Unfavourable | 2,21 |  |
| Chemokine inhibitor | LEU* | 105 | SER | WA2022 | Stabilising | Unfavourable | 0,33 | 0,33 |

We have taken under consideration the impact of the single missense mutation on the stability of the protein and how the distribution of main torsion angles φ and ψ influence the amino acid ability to three-dimensionally interact with the previous and next amino acid, reported in terms of predicted ΔΔG (kcal/mol) for the single missense mutation and in terms of overall predicted ΔΔG (kcal/mol) for the entire protein (Table 2)(supplementary figures 1-9). In particular the MPXV 2022 missense mutation have shown a Predicted ΔΔG of -0,57 for the mutation Y221H on the Immunomodulator A46 protein; a Predicted ΔΔG of 0,58 and -0,27 respectively for the mutation F54S and N313T on the TNFr protein; a Predicted ΔΔG of 2,21 for the mutation L307S on the Large Transcript protein 3 protein; a Predicted ΔΔG of 0,33 for the mutation L105S on the Chemokine Inhibitor protein and a Predicted ΔΔG of 0,99 on the mutation N266D of the Ankyrin like protein.

## Discussion

MPXV encodes homologs of mammalian cytokine and chemokine receptors that can dampen, evade, or subvert the host immune response and influence the outcome of infection (18). In our study we have found in all the 2022 MPXV isolates the presence of 5 missense mutations that were not previously reported in any other MPXV sequence. Most of these mutations appear to be likely able to affect metabolically and immunological factors, crucial in host-pathogen interaction. such as the Immunomodulator A46, the TNF receptor, the Ankyrin-like protein, the chemokine Binding protein and the Large Transcript protein (19). In the absence of direct experimental assessment, it is unclear whether and to what extent these missense mutations are affecting the function of the mutated proteins. Nonetheless, some of them have shown non-neglectable changes in energetic parameters which suggest that a functional impact of the mutation is indeed possible. Most of the mutations were found to be protein-stabilizing with an overall predicted positive ΔΔG (kcal/mol). One exception is the mutation Y221H on the A46 protein. This is a member of the poxviral Bcl-2-like protein family that inhibits the cellular innate immune response (20) as it has a destructive interaction with members of signalosome family MAL and MyD88 which play an essential role in host response to infection. A46 destabilization could decrease its ability to negatively interact with host factors above, thus favoring host immune response (21).

Another exception could be the TNFr protein. This MPXV factor belongs to a family of TNF receptor homologs acting as cytokine response modifiers able to dampen inflammatory response by the host, cellular destruction and inhibition of virus replication (22). Notably, the destabilizing value of TNFr mutation is low and appears to be compensated by an increased torsional ability thus shifting the overall predicted ΔΔG to a slight positive value, meaning the mutation might have a little impact on the host response. Regarding the Ankyrin-like protein, previous studies have reported that subtle variations directly influenced the cellular ubiquitination pathways and the anti-viral response coordinated by NF-κB. (23). Moreover, another stabilizing mutation has been found on the chemokine binding protein, this protein is well-known for its ability to interfere with host immune surveillance processes. (24).

The stabilizing mutation on the Transcription factor IIB (TFIIB) could potentially facilitate viral replication favoring the transcription initiation interacting with RNA polymerase II promoters. (25).

Both the stabilizing and the destabilizing mutations (particularly, the A46 mutation) may potentially influence MPXV-host interaction but increase viral fitness and make interhuman transmission easier. Indeed, in previous outbreaks of West African Clade the interhuman transmission has been reported to be less efficient than that observed with the isolates belonging to the (26) Central African Clade (27). We particularly speculate about a potential influence of some of the above mutations (in particular, those pertaining to A46 and TNFr) in restricting the inflammatory host response and disease symptomatology. These mutations appear to justify the paucisymptomatic clinical picture and high skin-mucosal viral tropism of the current MPXV outbreaks cases which systemic symptoms were mild, lacking even fever, asthenia or myalgias in most cases with high number of skin lesions that where mostly present in the contact area [23,24]. This inference does obviously need further direct investigations especially considering that, for Ubiquitin, Large Transcript and Ankyrin-like proteins, the homology modeling has shown a low sequence identity percentage.

Altogether, our results highlight how epidemic outbreaks that have occurred in recent years did not come from a strain that is accumulating mutations and that is circulating among humans but more likely from mutations occurring during different epizootic outbreaks due to Western African strain. The mutations usually occur on proteins responsible for immunomodulation, virus-host interaction and viral replication and tend to stabilize the protein structure.

Moreover, we can speculate that the progressive stabilizing mutations are the mechanism that led to the emergence of a highly adapted virus that is now causing a mild infection with higher mucosal tropism and sustained human-to-human transmission. In fact, patients appear to have a higher number of lesions in the contact area and no major systemic symptoms and when described are ascribed to mild symptoms such as myalgia, asthenia, and fever.

Even though MPX virus outbreaks rarely occur, partially due to its way of transmission, the molecular surveillance of this virus is crucial in order to prevent future outbreaks or mutations potentially inducing pathophysiological changes or broadening routes of transmission. In this article we have found and analyzed 5 new mutations that occurred during the 2022 MPXV outbreak, involved in host-human interaction modified in a positive or negative way the immunomodulation and replication pathways that are crucial for viral fitness. Given that, confirming these results and monitoring the possible emergence of new mutations will be essential to prevent future outbreaks potentially dangerous for public health.

## Acknowledgment

The authors thank the family of Prof. Giuseppe De Feo for the support. M.G. is funded by PON “Ricerca e Innovazione” 2014-2020.

